# NEURAL ENCODING OF FELT AND IMAGINED TOUCH WITHIN HUMAN POSTERIOR PARIETAL CORTEX

**DOI:** 10.1101/2020.07.27.223636

**Authors:** Srinivas Chivukula, Carey Zhang, Tyson Aflalo, Matiar Jafari, Kelsie Pejsa, Nader Pouratian, Richard A. Andersen

## Abstract

In the human posterior parietal cortex (PPC), single units encode high-dimensional information with *partially mixed* representations that enable small populations of neurons to encode many variables relevant to movement planning, execution, cognition, and perception. Here we test whether a PPC neuronal population previously demonstrated to encode visual and motor information is similarly selective in the somatosensory domain. We recorded from 1423 neurons within the PPC of a human clinical trial participant during objective touch presentation and during tactile imagery. Neurons encoded experienced touch with bilateral receptive fields, organized by body part, and covered all tested regions. Tactile imagery evoked body part specific responses that shared a neural substrate with experienced touch. Our results are the first neuron level evidence of touch encoding in human PPC and its cognitive engagement during tactile imagery which may reflect semantic processing, sensory anticipation, and imagined touch.

## INTRODUCTION

The posterior parietal cortex (PPC) is critical to integrating sensory information into motor plans and monitoring ongoing movement (1, 2). In recent studies, we identified at the level of single neurons, evidence that the human PPC encodes a wealth of movement related information including movement plans and trajectories, as expected, but also other variables such as cognitive motor imagery, action semantics, observed actions, and memory (3-7). This richness of representation is made possible through a *partially mixed* encoding in which single neurons represent multiple variables, allowing a relatively small neuronal population (recorded through a small 4×4 mmm implanted microelectrode array) to provide many movement related signals (6). Here, we explore whether this neuronal population also encodes the somatosensory domain, given its often intimate association with movement planning, initiation, and execution.

Multiple lines of evidence support somatosensory representations within the PPC. In non-human primates (NHPs), neurophysiological studies demonstrate somatosensory processing in cell populations around the intraparietal sulcus (IPS), where they are thought to play a role in higher-level cognition and perception (8-16). Examples include monitoring of limb configuration (through convergent visual and proprioceptive information) and sensing the space around the body (peripersonal space; convergent visual and tactile information) (8-11). In humans, lesion and neuroimaging studies support similar representations (17-19). Moreover, functional neuroimaging studies in humans demonstrate that experienced, observed, and imagined touch activate overlapping regions of the PPC, suggesting its role in a multisensory, cognitive processing of touch (20-25). While a sizeable body of literature has developed around somatosensory representations within the PPC, several fundamental questions remain. At the single neuron level, how are receptive fields to touch encoded? If bilateral information is represented, are the two sides discriminable? To what extent are cognitive touch representations activated during imagined touch encoded within the same neuronal populations? Does activity evoked during tactile imagery share a neural substrate with experienced touch?

In a unique opportunity, we investigated touch processing at the level of single neurons in a tetraplegic human subject recorded with an electrode array implanted in the left PPC for an ongoing brain machine interface (BMI) clinical trial. In previous work, we have referred to the implant area as the anterior intraparietal cortex, a region functionally defined in NHPs (3-6, 26). Here we will refer to the recording site as the postcentral-intraparietal area (PC-IP), acknowledging that further work is necessary to definitively characterize homologies between human and NHP anatomy. We recorded from a total of 1423 single neurons during the presentation of objective touch and during imagined touch to sensate dermatomes above the level of the participant’s injury. We found that human PC-IP neurons encoded experienced touch at short latency (∼100 ms) with bilateral receptive fields, covering all tested, sensate regions within the head, face, neck, and shoulders. Tactile imagery evoked body part specific responses that shared a neural substrate with experienced touch. Our results demonstrate for the first time, a high-fidelity, reproducible encoding of touch that can partially be reactivated during tactile imagery in a body part specific manner. The latter represents a novel finding, thus far untestable in NHP models, and suggests PPC involvement in the cognitive processing of touch.

## RESULTS

We recorded from 1423 well isolated and multi-unit neurons in the PC-IP (left-hemisphere) of a high cervical (level three to four; C3/4) spinal cord injured, tetraplegic human participant over 14 recording sessions at approximately one-week intervals (on average, 101.64 ± 7.22 neurons recorded simultaneously). Recordings were split across four tasks, designed to probe basic properties of the neuronal population relating to both experienced and imagined touch.

### PC-IP neurons encode bilateral tactile receptive fields

We first examined the hypothesis that PC-IP neurons encode tactile receptive fields to dermatomes above the level of the participant’s spinal cord injury (SCI). Tactile stimuli were delivered as rubbing motions at approximately 1Hz, for 3 seconds. The subject was asked to keep her eyes closed to eliminate neural responses arising from visual input. Tactile stimuli were presented to bilateral axial (forehead, vertex, cheek, neck, back) and truncal (shoulder) body parts to determine the extent of body coverage of any tactile representations among PC-IP neurons. As controls, touch was also presented to the bilateral hands (insensate regions below the level of SCI), and a null condition was included (with no stimulus delivered), to verify that touch related neural responses observed were not random.

A significant fraction of the neuronal population encoded touch to each of the tested body parts with preserved somatosensation (**Figure 1A**, *p*<0.05, with false discovery rate (FDR) correction). These results are consistent with bilateral encoding as the sensory fields included both left and right sides of the body. No significant modulation was seen in response to stimuli delivered to the hands, or in the null condition. Across the population, contralateral stimulation can be better discriminated (through a discriminability index, DI) from baseline activity (**Figure 1B**, *p*<0.05, sign test). Many neurons demonstrated an exclusive activation for touch to a single body part, although a substantial fraction of the population also demonstrated activation to multiple body parts (shown as a bar plot in **Figure 1C**). Sample neuronal responses of touch to various sites are shown in **Figure 1D**.

**Figure 1.**
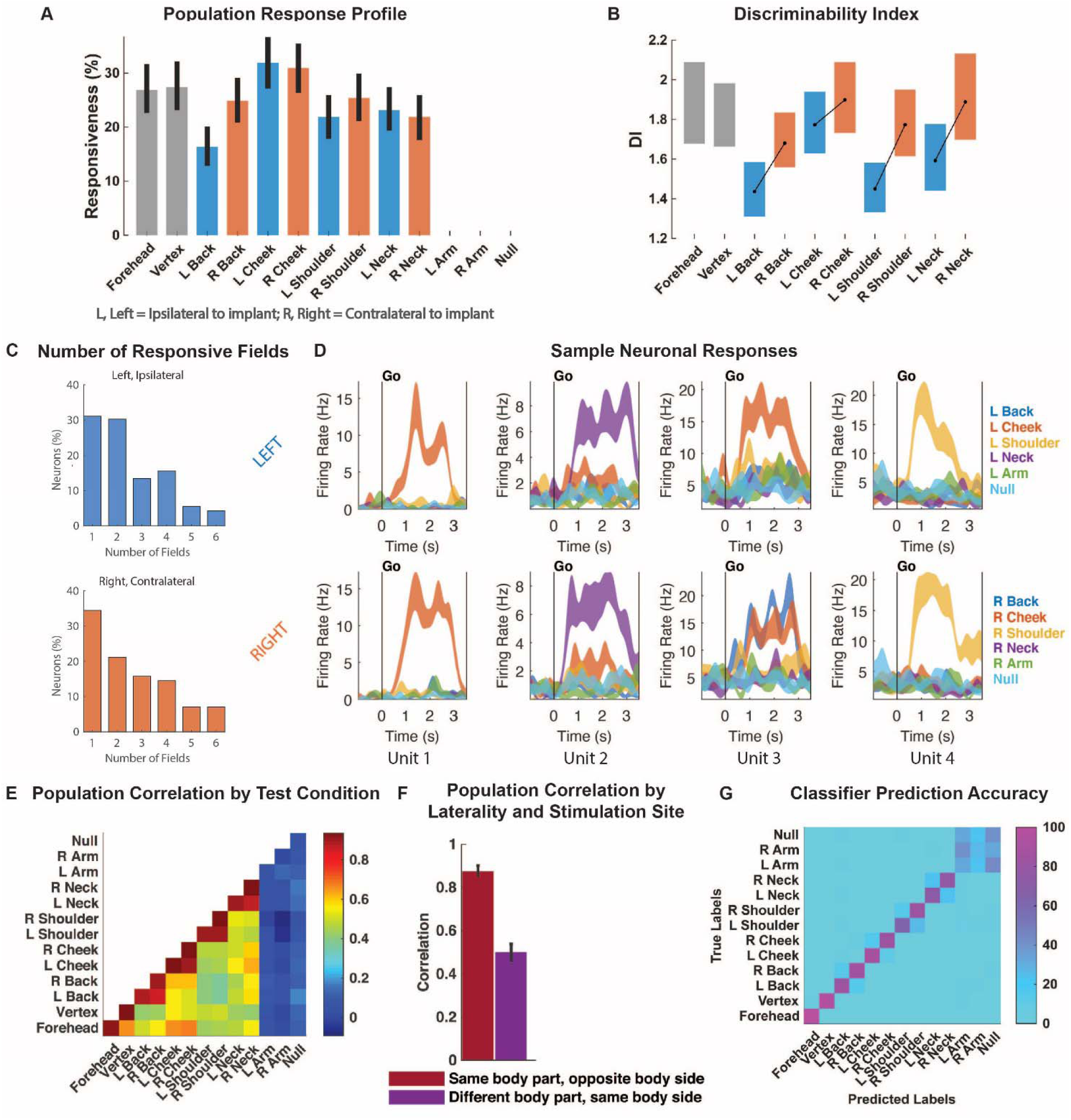
PC-IP encodes tactile stimuli. **A**, Percent of the PC-IP neuronal population that demonstrated significant modulation (tuning) to each tested stimulation site (bootstrap 95% CI, *p*<0.05, FDR corrected). **B**, Depth of neuronal modulation from baseline measured by discriminability index (DI, shown as 95% CI, see Methods) at each stimulation site. **C**, Bar plot of the number of fields to which the PC-IP neuronal population (expressed as percentage) significantly responded shown. **D**, Representative neuronal responses illustrating body part selectivity. Each panel shows the firing rate (mean ± SEM) through time. Each column illustrates the responses of the same unit to the left (top) and right (bottom) body side. **E**, Neuronal population correlation demonstrating the relation in encoding structure between test conditions. **F**, Population correlation for neuronal representations of touch to identical body parts on opposite body sides, compared to that for touch to different body parts on the same body side, for all tested sensate stimulation sites (mean with 95% CI). **G**, Confusion matrix of the cross-validated classification accuracy (as percentage) for predicting body parts from population neural data. The matrix is an average of the confusion matrices computed for each recording day individually.

### PC-IP population responses demonstrate spatial preference for body part

A neuron that encodes touch to a particular body-part could 1) be entirely specific to that body part only, 2) may respond to others, but show proclivity for touch to an alternate, ipsilateral field, 3) may prefer the corresponding, contralateral receptive field, or, 4) may show random patterns of activation to other sensory fields. We performed a correlation analysis of the population responses across each stimulation condition. The results are shown in **Figure 1E and Figure 1F**. The population responses to stimulation on the right and left sides of the body were highly correlated (**Figure 1E**), indicating that the pattern of activation across the neuronal population for a body segment is largely equivalent across the left and right sides of the body. For example, the population correlation between the left and right neck is similar to the cross-validated correlation of the left neck to itself. We note that response patterns to non-identical body parts (on either body side) are non-zero, suggesting a shared response to the simple presence of a stimulus (or potentially the precise type of stimulus, e.g. rubbing motions), independent of the precise location of the stimulus. A direct comparison of population correlation between matching body parts of the right and left side with population correlations within a body side is shown in **Figure 1F**.

We found that population neural activity can be used to accurately classify body parts above the level of injury, including differentiation of the body side (**Figure 1G**). However, in line with the correlation analysis, incorrect classifications tended to be for the matching body part on the opposite body side. Tactile stimuli to insensate hand regions was frequently confused with the null condition, consistent with the lack of any meaningful neural selectivity for these control conditions.

We examined bilateral coding at the level of individual neurons. We first investigated whether neurons selective for body parts on the contralateral side were also selective for body parts on the ipsilateral side. For each neuron we computed a linear model that described firing rate as a function of response to the stimulated body part, independently for the contralateral and ipsilateral body sides. For each linear model, we computed a cross-validated coefficient of determination (R^2^_within_) to measure the strength of neuronal selectivity for each body side. The R^2^_within_ values for the left and right for each neuron are plotted against each other in a scatter plot in **Figure 2A.** Most points cluster around the identity line, indicating that units highly selective for body parts on the left were also highly selective for the right side. This bilateralism is also reflected in the specificity plot shown in **Figure 2B**, in which it is evident that for most units the strength of selectivity was comparable across the ipsilateral and contralateral sides, with a slight bias for the contralateral side (*p*=0.04, sign test). This bias is consistent with the greater discriminability for touch to the contralateral body than to the ipsilateral body seen in **Figure 1B**.

**Figure 2.**
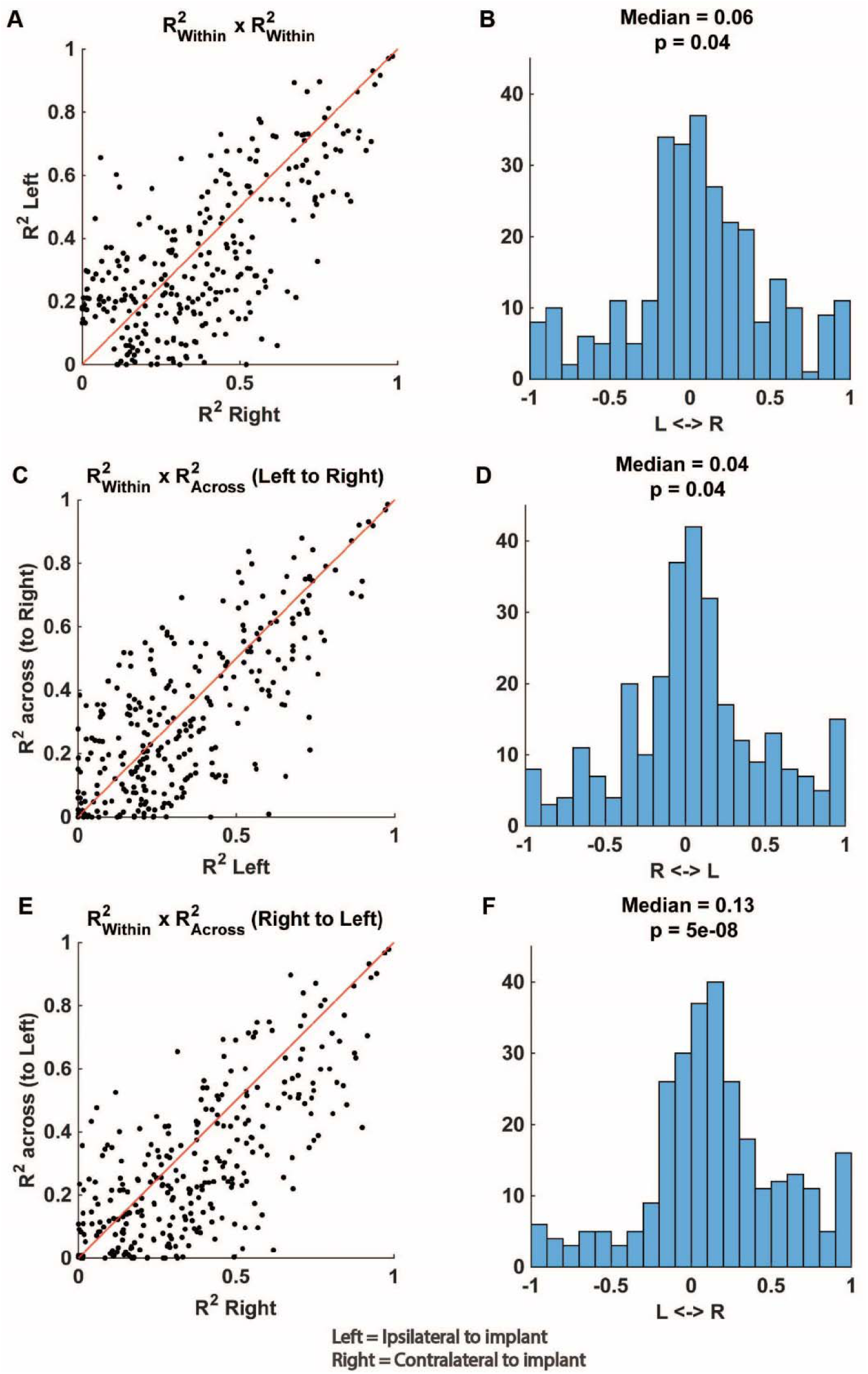
Touch responses are predominately bilateral. **A**, Scatter plot showing the strength of single unit selectivity (measured by cross-validated R^2^_within_, see Methods) to touch to the right side of the body compared to selectivity strength to touch to left side of the body. Along the identity line (in red), units have equal selectivity to the two body sides. **B**, Histogram of the side preferences of individual neurons. Side preference was quantified as the specificity index (see Methods) of each neuron, based on its response to touch for the right and for the left body sides (data from Figure 2 A). **C**, Scatter plot demonstrating how well selectivity patterns computed from neuronal responses to the left body side generalize to the right body side. Along the identity line (in red), units have the same pattern of selectivity for the right and left sides. **D**, Like 2B, but computing specificity index from data in 2 C. **E**, Like 2C, but now comparing right side generalization to the left side. **F**, Like 2 B, but computing specificity index from data in 2E. (R^2^, cross-validated coefficient of determination; L, left body, ipsilateral to implant; R, right body, contralateral to implant).

Next, we asked whether the precise patterns of responses observed for stimulation of the left body parts generalized to the right side, and vice versa. In other words, how does the population level similarity in coding for bilateral body parts manifest at the single unit level? To address this question, for each neuron, we trained a linear model to predict firing rate as a function of stimulated body part using contralateral data and predicted the firing rate of the ipsilateral data (and vice versa). The ability to predict across body side was quantified as the R^2^_across_ and compared to the R^2^_within_ computed above (**Figure 2C** and **Figure 2E**). We found that R^2^_across_ and R^2^_within_ clustered around the identity line, indicating a high similarity in encoding between the two sides for corresponding receptive fields. Specificity plots are shown in **Figure 2D** and **Figure 2F.** The centered distributions around a specificity index of zero reflect that response structures for most units are very similar for corresponding fields, with a slight preference for the contralateral side (*p*=0.04 and *p*<0.05, sign test, respectively). Critically, these results demonstrate that nearly all recorded PC-IP neurons demonstrate bilateral coding of tactile receptive fields across their range of representation strength (quantified by R^2^_within_).

### Tactile responses occur at short latency to bilateral stimuli

We explored PC-IP population encoding and single unit response latencies to tactile stimulation on the contralateral and ipsilateral body sides. In a variation of the basic task paradigm, we used a capacitive touch sensing probe to acquire precise measurements of the latency in neuronal response from the time of tactile stimulation. We probed latency on the bilateral cheeks and shoulders. Again, as a control, we included both hands in the task design. We compared latencies between the two sides at both the level of the PC-IP neuronal population as well as at the single unit level (as detailed in Methods).

At the population level, we measured encoding latency as the time at which stimulated body parts were discriminable based on a sliding window classification analysis. Encoding latency was short for both body sides and was slightly shorter for contralateral (right) receptive fields (96 ms) than for ipsilateral (left) receptive fields (104 ms) although this difference was not statistically significant. Figure 3A shows the time course of classification accuracy across the duration of a trial (sliding window, 150ms Full-Width Half-Max (FWHM) Gaussian smoothing kernel stepped at 50 ms). In figure 3B, the shaded region of 3A is expanded and shown with the minimal smoothing used for latency estimation (sliding window, 20 ms FWHM stepped at 2 ms).

**Figure 3.**
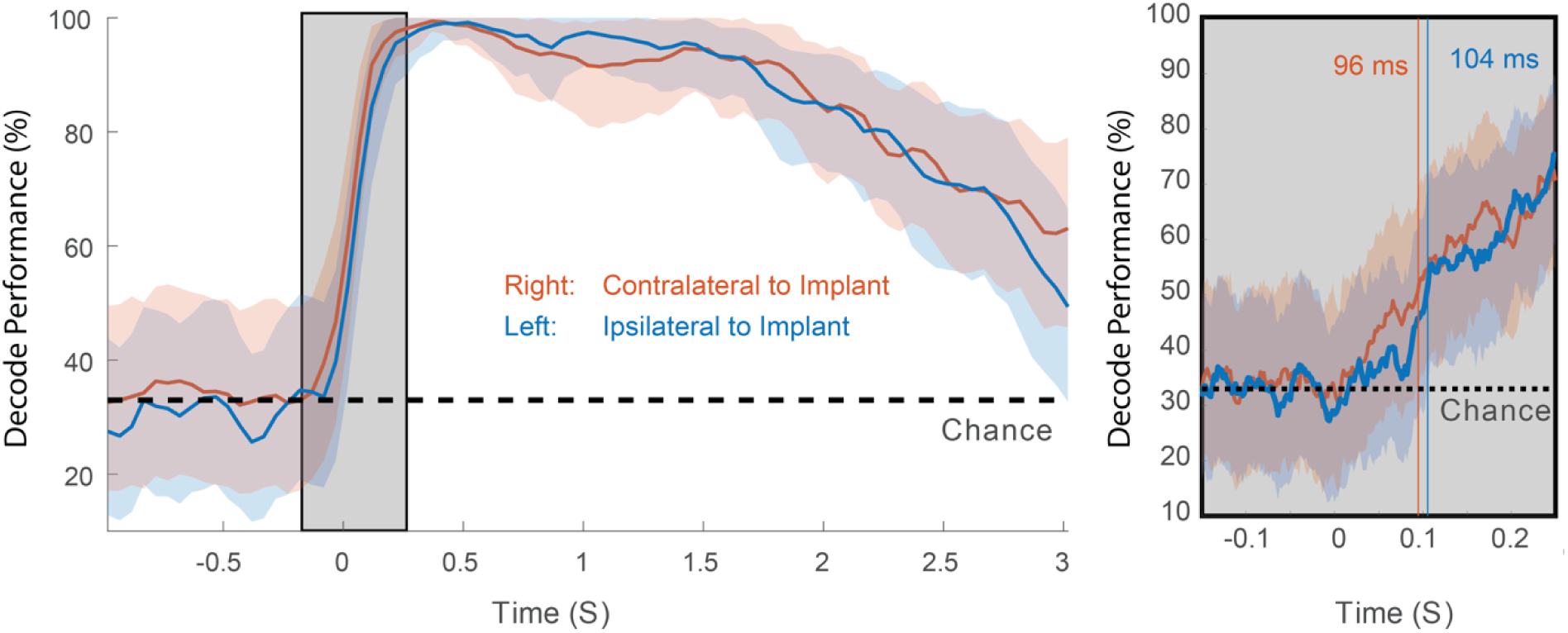
Tactile responses occur at short latency. Sliding window cross-validated classification accuracy (expressed as a percentage) aligned to time of contact (mean with 95% confidence interval). Classification accuracy was computed separately for the left (blue; ipsilateral to implant) and the right body sides (orange; contralateral to implant) and is shown as a function of time (50 ms step size with 150 ms smoothing). Chance accuracy is shown by the dashed horizontal black line. The boxed region is shown expanded on the right. In this panel, classification accuracy is shown at higher temporal resolution (2 ms step size with 20 ms smoothing). The vertical lines are color coded to indicate the decode latency (see Methods) for left and right body sides.

### Neuronal receptive fields to tactile stimuli on the cheek are spatially localized

We explored the receptive field structure within a body part at finer spatial precision to begin to characterize the sizes and shapes of receptive fields. We used a paintbrush to stimulate each of nine points equidistantly spaced along the participant’s cheek and neck, two centimeters apart, as shown in **Figure 4A**. A significant fraction of the neuronal population responded to tactile stimulation at each point, although the fractional responsiveness of the PC-IP population appears greater to stimulation points on the cheek than on the neck (**Figure 4B**). The strength of modulation from baseline, measured by the discriminability index, demonstrated a similar trend (**Figure 4C**). These findings are consistent with the results of **Figure 1A**, although all values are somewhat depressed, likely due to the gentler and spatially localized sensory stimuli. At the population level, responses demonstrate spatially structured receptive fields with stimulation of neighboring locations eliciting more similar activity than stimulation of distance locations (**Figure 4D**). Sample neuronal responses showing response as a function of stimulation site are shown in **Figure 4E**. Most neurons preferred a single stimulation site and demonstrated progressively less activity with increasing distance from that site.

**Figure 4.**
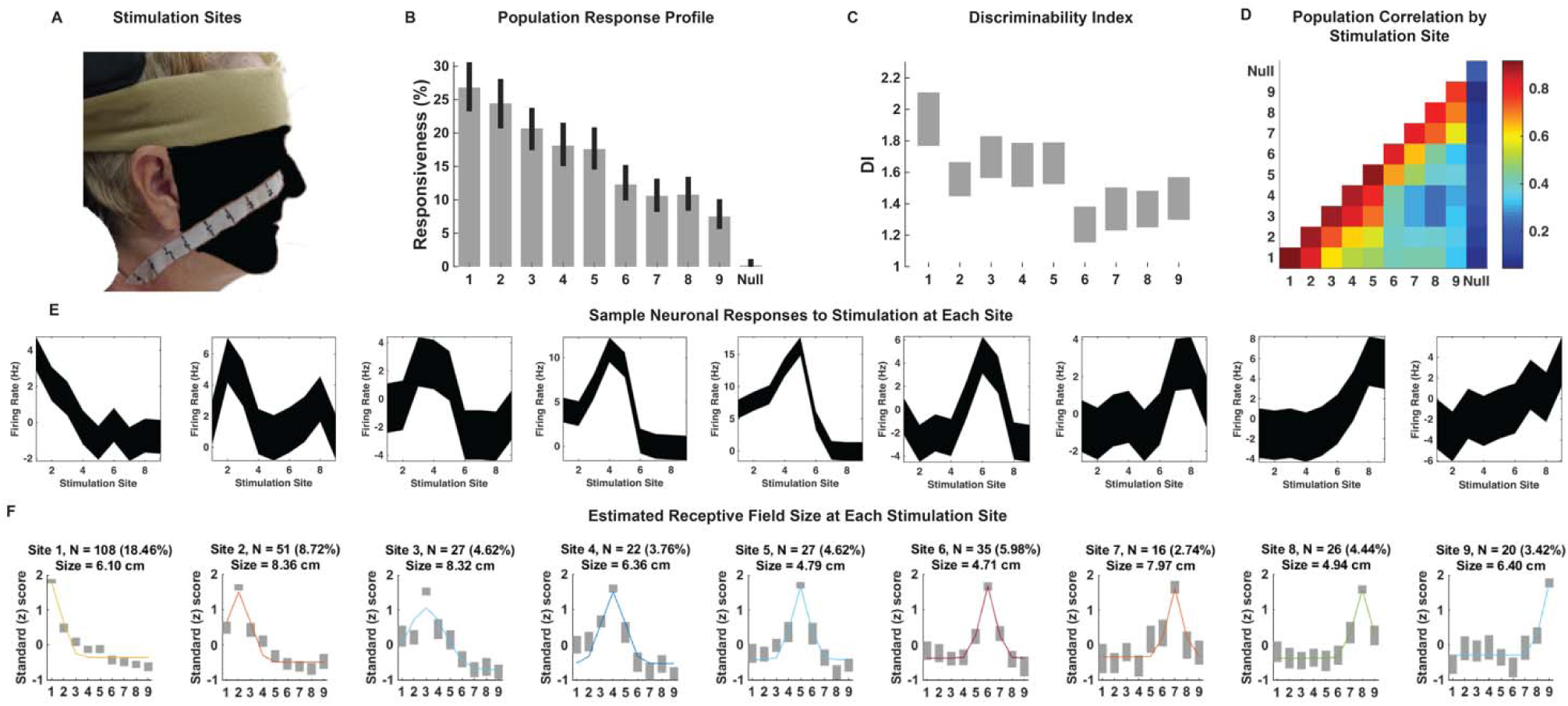
Receptive fields are spatially localized. **A**, Location of stimulation points along the study subject’s face and neck. Photo credit: Tyson Aflalo, California Institute of Technology. The face is masked to obscure identity per publisher’s request. **B**, Percent of the neuronal population significantly modulated to each stimulation point (mean with 95% CI, *p*<0.05, FDR corrected). **C**, Depth of neuronal modulation from baseline measured by discriminability index (DI, see Methods) at each stimulation site. The bar height represents the 95% CI around the mean value (midpoint). **D**, Population correlation matrix depicting the extent of similarity in encoding structure among the neuronal responses to touch at each stimulation point. **E**, Representative neuronal responses to stimulation at each site for example single units. **F**, Estimated receptive field size at each stimulation site. Well isolated neurons with a preferred response to touch at each stimulation site were selected, and their response to touch across stimulation sites modeled as a Gaussian function. The number of units included in the computation of this Gaussian curve is depicted above each subplot, including what percentage of the PC-IP neural population this comprised. The full width at half maximum of the Gaussian was used to estimate the size of the receptive field at each site, shown above each subplot. In each subplot, the x-axis indicates the stimulation site. The y-axis is a standard (z) score, representing how many standard deviations the mean spiking activity for neurons at each stimulation site was from the mean activity for that group of neurons at all sites together.

To estimate the average size of neuronal receptive fields (see Methods for details), we first identified all neurons demonstrating significant differential spatial representation of touch. For these units, categorized by their site of preferred (peak) response, we fit the standard deviation of a Gaussian function centered on the peak response to estimate the tuning width. **Figure 4F** shows the averaged responses for each stimulation site. The full width at half maximum (FWHM) of the neuronal receptive fields ranged from 4.79 cm to 8.36 cm and spanned, on average, between two and four stimulation sites.

### Tactile imagery evokes body part specific responses congruent with objective touch

Is the PC-IP recruited during tactile imagery? And if so, how might evoked neural responses compare to those arising from objective touch? To address these questions, we analyzed population activity elicited during a cue-delay-go tactile imagery task and compared the neural activity to that from objective touch to matching body parts recorded during interleaved trials. For imagery, the participant was instructed to imagine touch to the right (contralateral) cheek, shoulder, or hand with the same qualities as the objective touch stimuli the participant experienced during interleaved trials. A null condition was included as a baseline to measure neural activity without an objective or an imagined touch.

As with findings for objective touch, neuronal responses elicited by tactile imagery following the go cue (during the imagery phase or epoch) were discriminably encoded (**Figure 5A**, cross-validated accuracy=92%). When the null condition was excluded from this analysis, the prediction accuracy was higher, approaching 98%. High decode accuracy is consistent with the participant’s compliance with task instructions and implies that tactile imagery can elicit selective neural responses.

**Figure 5.**
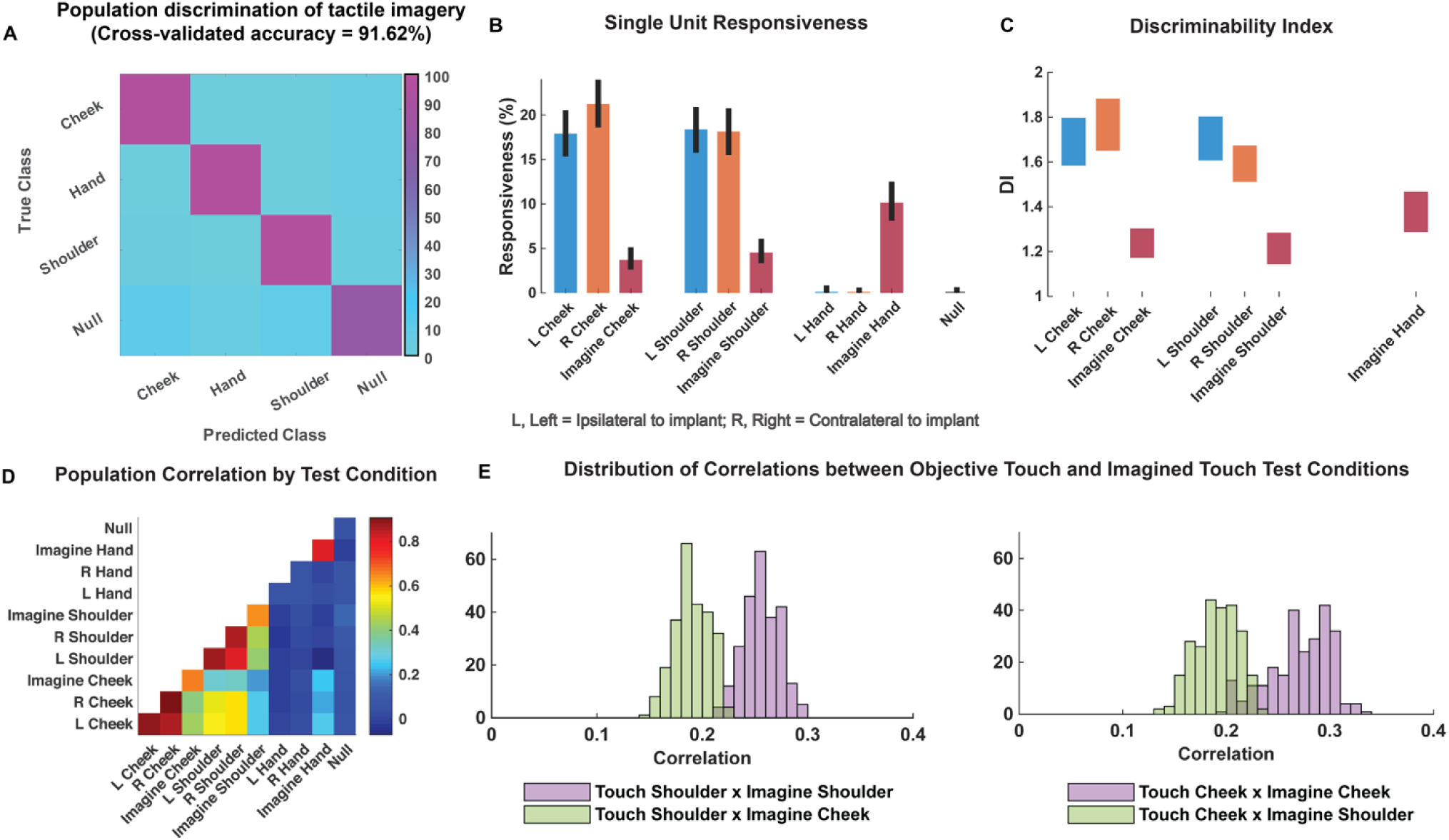
PC-IP neurons encode body part specific responses during tactile imagery. **A**, Average classification confusion matrix across recording sessions for body parts during tactile imagery and the baseline (null) condition. Colors represent prediction accuracy as a percentage, as in the scale. **B**, Number of neurons (as percentage) with significant modulation from baseline (bootstrap 95% CI, *p* <0.05, FDR corrected) split by test condition. **C**, Depth of neuronal modulation from baseline measured by discriminability index (DI, shown as 95% CI, see Methods) at each stimulation site. **D**, Population correlation matrix depicting similarity of the population response between all test conditions. **E**, Distribution of correlations between objective shoulder (left) and cheek (right) touch and imagined cheek/shoulder touches, with the distributions computed over different splits of the data (see Methods).

Consistent with previous results, a significant fraction of neurons encoded objective touch to the cheeks and shoulders but not to the hands. In comparison, a smaller fraction of the neuronal population was responsive to the cheek and shoulder during imagery of tactile stimuli (**Figure 5B)**. Of note, a significant number of neurons responded to imagined touch to the hand, despite the hand being clinically insensate in the study participant (and despite objective touch to the hand not eliciting neuronal activation). Discriminability from baseline neural activity, measured by the discriminability index, demonstrated a similar trend to the single unit responsiveness profile (**Figure 5C**).

We used the population correlation measure to compare population level neural activity across conditions (**Figure 5D**). Neural activity during tactile imagery shared a neural substrate with responses evoked by objective touch: representations for imagined touch and for experienced touch were more similar for matching body parts fields than for mismatched body parts (**Figure 5E**, permutation shuffle test *p*<0.05).

### Dynamic evolution of population coding between task epochs suggests additional cognitive processes

The analyses above were restricted to the mean neuronal activity following the go cue (e.g. during objective touch or during imagery) to allow a direct comparison with results reported for the previous paradigms. We now expand this analysis. During the tactile imagery task, the participant heard a verbal cue specifying a body part (verbal cue = “cheek,” “hand,” or “shoulder”) followed approximately 1.5 seconds later by a beep instructing the participant to imagine the stimulus at the cued body part on the right side of the body. This cue-delay paradigm is standard in the motor physiology literature and is used to dissociate planning from motor execution related neural activity (3, 27-29). In our case, the cue-delay was unique to the tactile imagery condition. We utilized the cue-delay task to begin to dissociate in time whether neural activity during tactile imagery is consistent with the neural correlate of imagined touch.

To leverage the benefits of the cue-delay paradigm, we performed a dynamic classification analysis (500ms windows, stepped at 100ms, see Methods). Results are shown as a matrix (see **Figure 6**). The diagonal elements represent the cross-validated prediction accuracy for a specific time window. The off-diagonal elements represent how well the classifier generalizes to alternate time windows. Each row can be interpreted as quantifying how well decision boundaries established for the diagonal time windows generalize to other time windows. This analysis allows us to measure when the neuronal population represents the different body parts (the diagonal) and whether population coding is similar or distinct during the task phases (the off-diagonal). We are interested in two main phases of the task, the early portion comprised of the cue and delay (cue-delay), and the later portion when the participant is actively imagining the stimulus (go/imagery). **Figure 6A** schematically illustrates examples of possible results. The examples are meant to be illustrative but are not an exhaustive list of possibilities. The population may be selective exclusively during the imagery phase, during the cue-delay and imagery phases but with distinct population coding, during the cue-delay and imagery phases with identical coding, or during the cue-delay and imagery phases with partially shared and partially distinct coding. Each pattern would suggest a different interpretation of various forms of cognitive processing that may be involved in tactile imagery (see Discussion).

**Figure 6.**
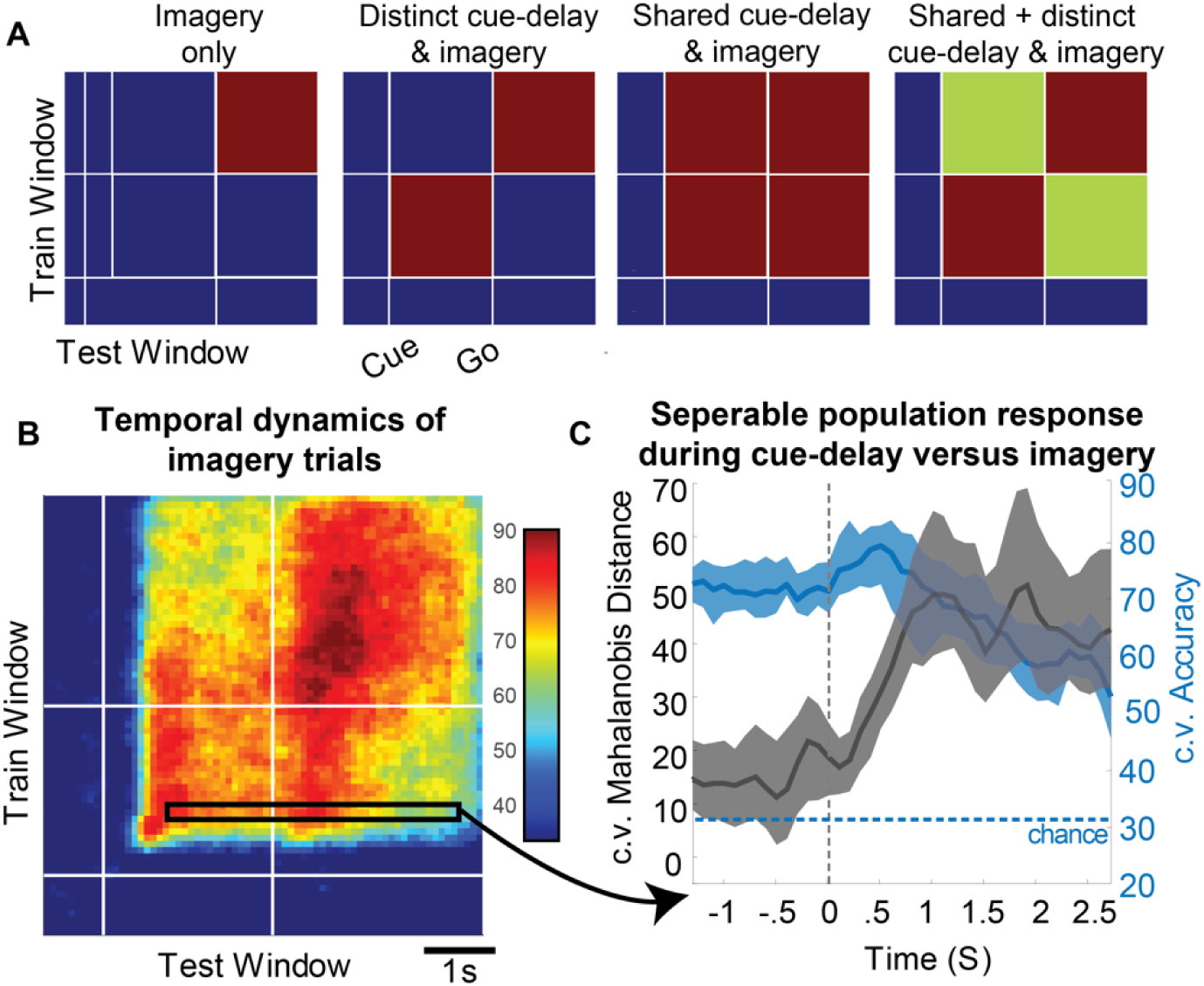
Shared and distinct coding of body parts during cue-delay and imagery epochs. **A**, Schematic illustrating possible dynamic classification patterns over epochs of the tactile imagery task. In each panel, the window used for classifier training is along the y-axis, and the window used for classifier testing is along the x-axis. The start of the auditory cue (marking the onset of the cue-delay epoch) and the beep (marking the go signal for the imagery epoch) are shown as solid white lines, labeled “Cue” and “Go.” **B**, Dynamic classification analysis results for the imagined touch test conditions with conventions as in 6A. The colors represent prediction accuracy values (as percentage) as in the scale. **C**, Illustration of distinct and shared neuronal responses between the cue/delay and imagery epochs for boxed window of of 6 B. Shared response illustrated with classification generalization accuracy (blue, mean with 95% confidence interval computed across sessions). Distinct response illustrated with cross-validated Mahalanobis distance (dark grey, mean with 95% confidence interval computed across sessions). The dashed vertical line marks onset of the imagery epoch. The dashed horizontal line marks chance classification accuracy. (c.v., cross-validated).

The results of our classification analysis (**Figure 6B**) are most consistent with body part (or sensory field) selectivity during both the cue-delay and imagery phases with partially shared and partially distinct populations coding of the body parts between phases. The shared component is evident in the significant generalization accuracy in the off-diagonal elements, a representative row of which is shown in **Figure 6C** (blue portion) where cross-validated accuracy generalizes from approximately 70% within the cue-delay phase to approximately 60% during the imagery phase. The distinct population activity between phases is best revealed by a cross-validated Mahalanobis distance as it provides a sensitive measure of change which is masked by the discretization process of classification (see Methods). The findings demonstrate a significant change between the activity patterns in the cue-delay and the imagery epochs (**Figure 6C**, black).

To further clarify the properties of individual units, we conducted a dynamic classification analysis for each recorded unit. This resulted in the same matrices described above, but now each matrix represents how information coding evolves for a single unit. Two complimentary analyses were then performed. In the first, a cluster analysis was performed on the resulting matrices (**Figure 7A**). Three clusters were identified that most parsimoniously accounted for observed activity patterns (Bayesian information criteria test for optimal number of clusters). Clusters roughly corresponded to temporal profiles with selectivity during the cue-delay and imagery phases with similar coding (30%), and units exclusive to the imagery epoch (33%) or the cue-delay epoch (37%). In the second analysis, time resolved classification data were analyzed using a principle components analysis (PCA), the first three principle components of which are shown in **Figure 7B**. A majority of variance (26%) is explained by units that are active during both epochs with similar coding.

**Figure 7.**
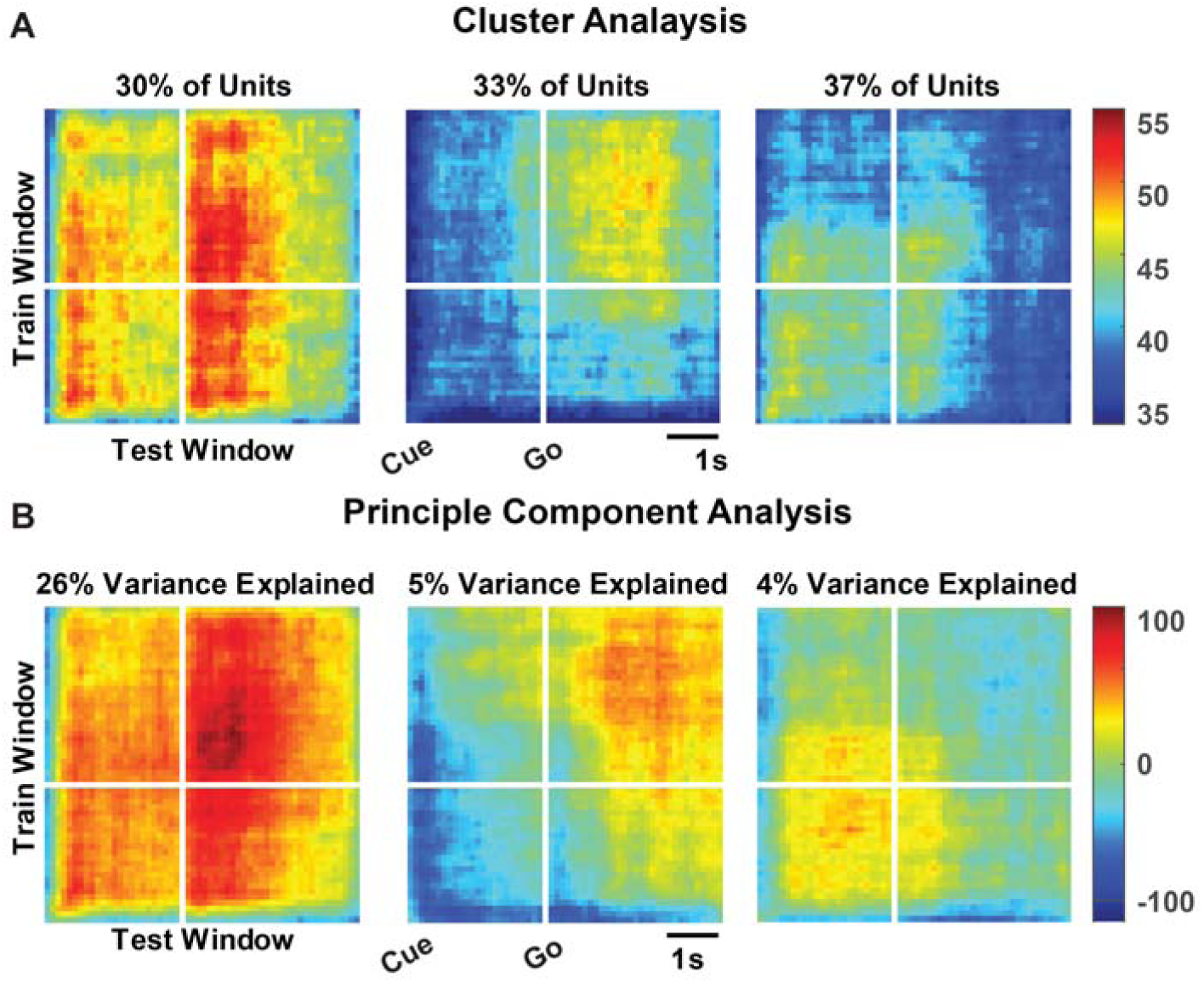
Diverse temporal dynamics in single units. **A**, Dynamic classification matrices were constructed separately for all selective units (see Methods). Resulting matrices were clustered to identify common temporal dynamics in individual units. Resulting clusters are visualized as the mean across all matrices assigned to the same cluster. The fraction of neurons (as a percentage of the selective neuronal population) assigned to each cluster is indicated. Plot conventions as in Figure 6. **B**, Principle components (PC) of the dynamic classification matrices of single unit activity (same data as 7 A) are shown, along with the fractional variance explained by each. Conventions as in 7 A except color PC weights.

### Cognitive processing during the cue-delay and imagery epochs of the tactile imagery task shares a neural substrate with that for objective touch

Finally, we look at how encoding patterns through time generalize between the tactile imagery and objective touch conditions. The dynamic classification analysis above was applied both within the imagery condition and across condition types (e.g., from imagined to experienced touch on the right side and vice versa; **Figure 8A**). We found significant generalization when training the classifier on either the cue-delay or the imagery phases of the imagery task and applying the classifier to the stimulus phase of the objective touch condition (**Figure 8A**, lower-left panel). This implies that cognitive processing prior to active imagery as well as during imagery share a neural substrate with experienced touch. A visualization of significant generalization from imagery to experienced touch for the boxed region of Figure 8A is shown graphically in **Figure 8B**. We did find an asymmetry in across-condition classification; training a classifier on experienced touch did not generalize well to the imagined touch condition (**Figure 8A**, upper-right panel). This asymmetry is likely a consequence of the analysis technique and may not be of physiological significance. **Figure 8C** illustrates the likely driver of the asymmetry using classifiers trained on the first principal component of the population response. The figure demonstrates how a decision boundary for relatively low signal-to-noise ratio (SNR) conditions will generalize to a higher SNR class, but not vice versa. Sample neuronal responses that help to understand single unit and population behavior are shown in **Figure 8D**.

**Figure 8.**
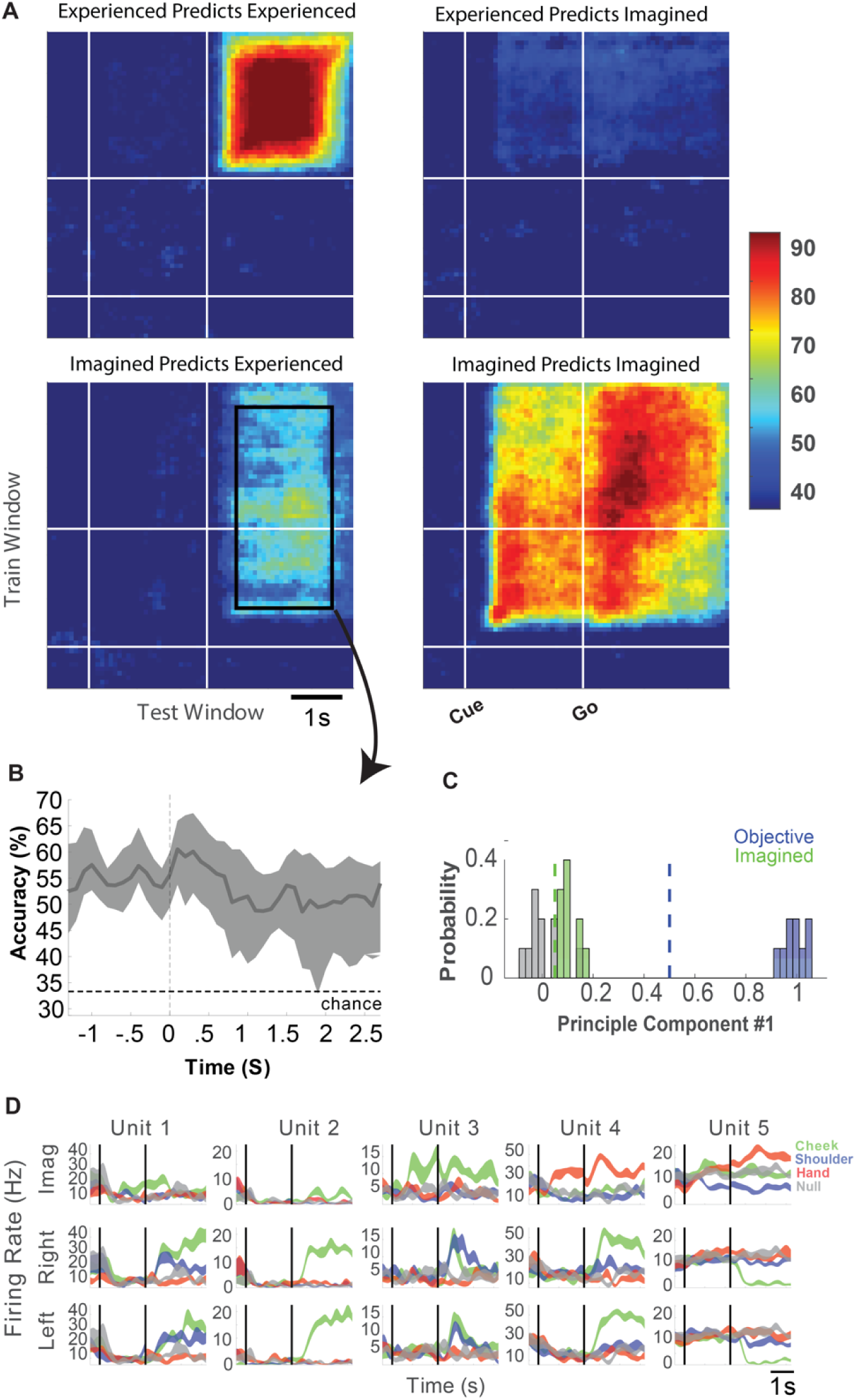
Cue-delay and imagery evoked neural activity shares a neural substrate with experienced touch. **A**, Within- and across-condition dynamic classification analysis demonstrating shared neural substrate between imagined and experienced tactile sensations. Upper-left and lower-right panels show within-format cross-validated accuracy for experienced and imagined sensations respectively. Upper-right and lower-left panels show generalization accuracy when predicting imagined responses based on experienced data and when predicting experienced responses based on imagined respectively. Conventions for each matrix are as in Figure 6. **B**, Generalization accuracy (mean with 95% CI) showing results for the boxed region in lower left panel of A. The dashed horizontal line marks chance accuracy. **C**, Illustration of how decision boundaries computed on data with different signal magnitudes can result in asymmetric generalization (see Methods). **D**, Representative neuronal responses illustrating selectivity during experienced and imagined sensations. Each panel shows the firing rate (in Hertz, mean ± SEM) through time (Vertical lines signal onset of cue/delay and go phases as labeled). Each column illustrates the responses of the same unit to tactile imagery of the right side (top), experienced sensations on the right side (middle), and experienced sensations on the left side (bottom) for matched sensory fields.

## DISCUSSION

We have previously reported that human PPC encodes many action variables in a high-dimensional and *partially mixed* representation (5-7). This architecture allows many parameters to be encoded by a small number of neurons, while still enabling meaningful relationships between variables to be preserved. Here we show that neurons recorded from the same electrode array in the same clinical trial participant are also selective for bilateral touch at short latency. Responses to objective touch are organized around body part, sharing population representations between the left and right side. Additionally, tactile imagery elicits body part specific responses that share a neural substrate with that for experienced touch. Furthermore, we found neural selectivity during active imagery as well as during the cue and delay epochs that precede imagery. The distinguishable population activity during these different phases indicates an encoding of multiple cognitive processes that may include semantic association, memory, sensory anticipation, or imagery per se.

### Experienced touch representation in human PC-IP is bilateral, body part centric and at short latency

Cortical processing of somatosensory information begins in the anterior portion of the parietal cortex (APC) within four cyto-architectonically defined areas termed BA 3a, 3b, 1 and 2 (30-32). Each of these four sub-regions represents primarily contralateral somatosensory information (33-40). Bilateral encoding in representation is thought to arise within the banks of the IPS, where the primary sensory regions blend into the PPC (38-41). Additionally, spatially localized and segregated sensory representations in neurons become progressively more integrated, and neuronal receptive fields become larger and more extensive, moving from the APC to the PPC (42-50).

We identified predominantly bilateral encoding of touch responses at short latency within the human PC-IP. Neuronal receptive fields were variable in shape and in extent of modulation from baseline, but demonstrated a preferred region of maximal response, with declining activity as a function of distance from the peak response. Further, many regions of the body were encoded within the small 4×4 mm cortical patch under the microelectrode array. Bilateral encoding of body parts (or receptive fields) in the PPC is consistent with findings in NHPs (35, 49, 51-54). We note, however, that most electrophysiological studies in NHPs examined tactile responses to the hands or forelimbs (8-12). Because these were unavailable for evaluation in our experiments (due to the study participant’s high cervical spinal cord injury level), we instead investigated and identified encoding of touch to bilateral receptive fields on the head, neck, and shoulders.

The latency to neuronal discrimination was short and consistent with bottom-up sensory processing of tactile stimuli. The latency was slightly shorter for contralateral than for ipsilateral stimuli, although the difference was small and not statistically significant. Contralateral touch also had marginally greater discriminability in neuronal modulation from baseline. Similar findings have previously been reported in animal studies (55, 56). For example, neurons in BA 2 and BA 5 in NHPs have bilateral receptive fields to visual and somatosensory stimuli, but with a contralateral limb bias (52, 56, 57). Support for a similar bilateral somatosensory response in human PPC is lacking but human functional neuroimaging studies have reported that blood oxygen level dependent (BOLD) signals are elevated bilaterally in the PPC in response to pointing movements of the fingers of either arm, albeit with a contralateral bias (58). The slight bias for contralateral tactile processing may be due to the different pathways by which information reaches these neurons from the two sides of the body. Thalamocortical afferents conduct primarily contralateral information to APC neurons, and possibly to PPC (and PC-IP) neurons as well (39, 40, 51-53, 55). Ipsilateral information, however, is believed to reach these neurons predominantly via cortico-cortical routes from the contralateral hemisphere (55, 56, 59).

### Tactile imagery dynamically invokes multiple cognitive processes in human PC-IP

Functional neuroimaging data in humans suggest that a network of brain regions is activated during both experienced and imagined touch. Involved brain regions include the PPC as well as portions of the insula (in particular, the posterior insula), the amygdala, and the bilateral temporal cortices (23, 60, 61). Human neuroimaging studies also support a role for the PPC in the interpretation of observed (not experienced) touch to others (20-22, 25, 62, 63). These observations collectively suggest that the PPC represents a node within a network of brain regions that support the shared processing of experienced, observed, and imagined touch.

In motor neurophysiology, neural activity related to planning and execution are dissociated in time by introducing a delay between the cue instructing movement, and the movement in response to the cue (27, 28). We have found that such distinctions between planning and execution are preserved during motor imagery experimental paradigms in tetraplegic individuals (3). Here, a similar paradigm allowed temporal dissociation in cognitive processing during tactile imagery. Single units demonstrated three dominant response profiles: 1) a shared selectivity pattern between the cue-delay and imagery epochs, consistent with cognitive engagement during all phases of the imagery task, 2) selectivity exclusively during the cue-delay epoch but not the imagery epoch, and 3) selectivity exclusively during the imagery epoch but not the cue-delay epoch. In a previous study, we found similarly heterogeneous responses during the cue, delay and imagery epochs for imagined hand grasp shapes (64). These single unit temporal selectivity profiles provide a basis for the population level findings of generalization in classification results between the cue-delay and the imagery epochs (**Figure 6B and Figure 6C**) but also a separation in neural state-space between these epochs (**Figure 6C**).

We acknowledge a limitation: while our task is well designed to identify dynamic engagement of multiple cognitive processes during tactile imagery, it is not adequate to precisely define the cognitive correlates of observed neural activity. This will be a subject of future investigation. A conjecture is that neural activity during the cue-delay and imagery epochs may reflect a combination of semantic processing of the verbal cue, sensory memory, sensory anticipation of a tactile stimulus, and imagery itself (4, 7, 65). An involvement of semantic processing is especially likely as we recently reported processing of read action verbs within the same neuronal population (7). The current findings would extend these results to the tactile domain and demonstrate neuronal selectivity for auditory cues (in addition to written text used in the previous study). It may be that the neurons involved in semantic processing within the PC-IP are shared whether for sensory (tactile) processing or motor planning. However, future work is necessary to better characterize the properties of these neurons as they relate to body part specificity, modality (sensory or motor) dependency, and other properties.

One concern with the use of all imagery experiments is that participant compliance cannot be externally validated. This raises the possibility that the participant is not performing the task or is performing the task in an unexpected manner. We think this is unlikely for three reasons. First, the subject by the time of this study was well versed in performing cue-delayed paradigms in the motor domain using both motor imagery and overt movements (e.g. 6). In Zhang and Aflalo et al. 2017, the participant’s performance of overt movements was perfect: the participant both performed the correct cued action and performed the action at the go cue (e.g. no movements began prior to the go cue as validated by measurements of electromyogram activity). Second, our current pattern of results that includes stable and accurate (near 100%) body part specific encoding within the cue-delay and imagery epochs, with a shift between epochs, is consistent with the participant performing the task as instructed. At a minimum, it is consistent with the participant’s performing two distinct cognitive operations during the two primary phases of the task with remarkable trial to trial consistency. Third, evidence for a shared neural substrate between experienced touch and the imagined touch conditions (discussed more below) indicates that selective responses during the imagery task are related to tactile cognition.

### Tactile imagery shares a neural substrate with experienced touch in human PC-IP

We found that experienced touch and the cognitive activity evoked by imagined touch shared a neural substrate within the PC-IP. Imagined touch to the cheek, for example, was more similar in representation to experienced touch to the cheek than to experienced touch to the shoulder, and vice versa. Interestingly, the overlapping neural representations between experienced touch and imagery were not limited to the stimulus phase (objective touch and imagery) itself, but also extended to the cue-delay phase of the imagery task. This overlap is consistent with our recent findings for shared neural representations between imagined and attempted actions, and for shared neural representations between observed actions and action verbs and is in line with our findings of a *partially mixed* architecture within PC-IP (5, 6, 66). These studies are all also consistent with views in which cognition recruits sensorimotor cortical regions (67-71). As with all passive neural recording studies, ours cannot establish a causal role for these neurons in tactile cognition. Understanding the unique contribution of PC-IP neurons within the larger network of brain regions engaged in cognitive touch processing remains to be explored. Nonetheless, our current results provide the first human single unit evidence of a shared neural substrate between tactile imagery and experienced touch.

A substantial fraction of the neuronal population activated in response to imagined touch to the hand, where no response to objective touch was seen (insensate in the study participant). This suggests that despite the lack of peripheral input from the hand due to the participant’s spinal cord injury, the brain maintains an internal representation of tactile sensations (72). The findings that intracortical microstimulation produces discernable tactile perceptions from insensate body regions adds definitive evidence for a maintained representation of somatosensory sensations after deafferentation (73, 74). These findings will prove useful for bidirectional neural prostheses.

### PC-IP and plasticity following spinal injury

The extent to which the human PPC reorganizes following SCI is unknown. Lesion studies in NHPs suggest that BA 3b and 3a, 1 and 2, show altered sensory maps following SCI, in a manner dependent on thalamic input from the afferent sensory pathways such as the dorsal column-medial lemniscus system (75). With mid-cervical lesions, for instance, there is an initial loss of BA 3b hand representations, and a slight expansion in face representation at approximately two years (75, 76). Although significant axonal sprouting has been demonstrated to occur at the site of deafferentation in the spinal cord, with increased projections to brainstem nuclei, the changes observed in the somatosensory cortex are significantly smaller (75, 76). Moreover, the reorganization in higher order somatosensory centers such as the secondary somatosensory cortex is even more restricted than in BA 3b (76). Against this background, we acknowledge that while additional work probing cortical reorganization following SCI is necessary to fully understand its electrophysiological consequences, our results within the current report provide insight into the maintenance of basic tactile processing within the human PC-IP, and PPC, after SCI.

## CONCLUSION

Multiple lines of evidence indicate a critical role for the human PPC in the integration of convergent multimodal sensory information to enable complex cognitive processing and motor control. To date, however, its processing of somatosensory information at the single neuron level has remained fundamentally unexplored. In the unique opportunity of a neuroprosthetic clinical trial, we examined the neural encoding of real and imagined touch within the human PC-IP. We found that local populations of PC-IP neurons within a 4×4 mm patch of cortex encode bilateral touch sensations to all tested sensory fields above the level of the participant’s injury at short latency. A significant fraction of PC-IP neurons encoded imagined touch with matching sensory fields to experienced touch. The activity in the delay period of the task, between cueing and imagining touch, may reflect cognitive processes including tactile semantics, sensory anticipation, as well as active imagery. Together, our results provide the first single unit evidence of touch processing within the human PC-IP and identify a putative substrate for the encoding of cognitive representations of touch, thus far untested in animal models.

## MATERIALS AND METHODS

### Study participant

The study participant, NS, is a 60-year-old tetraplegic female with a motor complete spinal cord injury (SCI) at cervical level C3-4 that she sustained approximately ten years prior to this report. She has intact motor and sensory function to the level of her bilateral deltoids. NS was implanted with two 96-channel Neuroport Utah electrode arrays (Blackrock Microsystems model numbers 4382 and 4383) six years post-injury, for an ongoing BMI clinical study. She consented to the surgical procedure as well as to the subsequent clinical studies after understanding their nature, objectives and potential risks. All procedures were approved by the California Institute of Technology, Casa Colina Hospital and Centers for Healthcare, and University of California, Los Angeles Internal Review Boards.

### Implant methodology and physiological recordings

The electrode implant methodology in NS has been previously published (3, 5, 6). One array was implanted at the junction of the left intraparietal sulcus with the left post-central sulcus in what we refer to as PC-IP. The other was implanted in the left superior parietal lobule (SPL). Implant locations were determined based on preoperative functional magnetic resonance imaging (fMRI). The participant performed imagined hand reaching and grasping movements during a functional MRI scan to localize limb and hand areas within this region. Following localization, a craniotomy was performed on August 26, 2014. The PC-IP electrode array was implanted over the hand/limb region of the PPC within the dominant (left) hemisphere, at Talairach coordinates [-36 lateral, 48 posterior, 53 superior]. In the weeks following implantation, it was found that the SPL implant did not function. Although this electrode array was not explanted, only data recorded from the PC-IP implant were used in this study.

### Experimental setup

All experimentation procedures were conducted at Casa Colina Hospital and Centers for Healthcare. Participant NS was seated in a motorized wheelchair in a well-lit room. Task procedures are presented in detail in the sections below. For most tasks, however, one experimenter stood directly behind the participant and was responsible for providing tactile stimuli to the participant. A 27-inch LCD monitor was positioned behind NS (visible to the experimenter but not to the subject) to cue the experimenter for the presentation of a stimulus. Cue presentation was controlled by the psychophysics toolbox (Brainard, 1997) for MATLAB (Mathworks) (77).

### Data collection and unit selection

Data were collected over a period of approximately eight months in the fourth year after NS was implanted. Study sessions were conducted between two and three times per week, lasting approximately one hour each. Neural activity recorded from the array was amplified, digitized, and sampled at 30 kHz from the electrodes using a Neuroport neural signal processor (NSP). The system has received food and drug administration (FDA) clearance for less than thirty days of recording. We received an investigational device exemption (IDE) from the FDA (IDE #G120096, G120287) to extend the implant duration for the purposes of the BMI clinical study. Putative neuron action potentials were detected at threshold crossings of −3.5 times the root-mean-square of the high-pass filtered (250 Hz) full bandwidth signal. Each individual waveform was made of 48 samples (1.6 ms) with 10 samples prior to triggering and 38 samples after. Single and multiunit activity was sorted using Gaussian mixture modeling on the first 3 principal components of the detected waveforms. The details of our sorting algorithm have been previously published by our group (6). Single units were pooled across recording sessions. Each recording was assumed to be independent and no assumptions were made about the same units being recorded on more than one study session. To minimize noise and low-firing effects in our analyses, we used as a selection criterion for units, a mean firing rate greater than 0.5 Hz and a signal to noise ratio (SNR) greater than 0.5. We defined SNR for the waveform shapes as the difference between their mean peak amplitude and the baseline amplitude, divided by the variability in the baseline.

For measurements of neural latency to stimulus response (please refer to the task descriptions below for more information), a custom capacitive probe was used to record the exact time of tactile stimulation. This probe was built using a Raspberry Pi 2B and Adafruit Capacitive Touch Hat (Adafruit product ID 2340). The digital output (a binary output for touch or no touch) was transmitted through a BNC cable into the NSP at an analog signal sampling rate of 2 kHz.

### Task procedures

We used several experimental paradigms to probe various features of experienced and imagined touch representations in the PC-IP. In each paradigm, the participant was instructed to keep her eyes closed. The basic task structure comprised three phases: Each trial began with the presentation of a cue to the experimenter (or an auditory cue in the tactile imagery condition, see specific task description below), 1.5 seconds in duration, indicating the stimulus (for example, touch NS’s left cheek). This was followed by a brief delay, 1 second in duration. Then written text appeared on the screen to signal the experimenter to present the instructed stimulus for 3 seconds (in the tactile imagery paradigm, a beep indicated the “go” signal for the participant). Exact time intervals varied depending on task. Trials were pseudorandomly interleaved; all conditions were necessarily required to be performed at least once before they were repeated. In tasks in which both left and right body sides (ipsilateral and contralateral to the implant, respectively) were tested, stimuli were delivered to one body side at a time.

#### Neural responsiveness to touch

This task variant explored neuronal responsiveness and selectivity to objective touch to body parts (receptive fields) with preserved somatosensory input (above the level of SCI). Body parts tested included the forehead, vertex of the head, left and right back of the head, left and right cheeks, left and right sides of the neck, and the dorsal surfaces of the left and right shoulders. As controls, the left and right hands (clinically insensate) and a null condition (no stimulus presentation) were also included. Objective touch stimuli were presented to each body part as finger rubs by the experimenter at approximately one per second. Stimuli to the left and right body sides were delivered on separate trials to evaluate each side independently. To ensure that any neural activity observed truly arose from experienced touch and not from observed touch or other stimuli, NS was instructed to close her eyes throughout the task. She additionally wore ear plugs to block auditory input. This task was performed on four separate days. On each day, ten trials per condition were conducted. In total, we recorded from 398 sorted units on four separate testing days.

#### Neural response latency

The purpose of this task was to determine the latency of neural response to objective touch for the left and right sides of the body. Tested regions included the left and right cheeks, the left and right shoulders, and as controls, the left and right hands (insensate). Objective touch stimuli were presented as in the task above. Instead of finger rubs, however, a capacitive touch probe was used to enable precise delineation of the actual time of contact (touch) before the onset of a neural response. This task was performed on eight separate days, with eight trials per condition in each run of the task. In total, we recorded from 838 sorted units.

#### Receptive field size

This task aimed to estimate the size of neuronal receptive fields to objective touch. Neural responses to nine equally spaced points were evaluated, two centimeters apart, along a straight line from NS’s right cheek to her neck (**Figure 4A**). Only the right side (contralateral) was tested in this task. The first of these nine points was on the cheek bone or the malar eminence, and the ninth point was on the neck as shown. In addition to the nine points, a null condition (no stimulus presentation) was also added. Stimuli were presented through a paintbrush gently brushed against each of the points, at a frequency of one brush per second. The paintbrush was employed to deliver spatially localized sensations without accompanying skin distortion that could mechanically stimulate nearby sensory fields. Data were recorded on six separate days. On each day, ten trials of each condition were tested. In total, we recorded from 585 sorted units.

#### Engagement during tactile imagery

This task was intended to establish whether PC-IP neurons are engaged by tactile imagery, and whether neural patterns evoked by cognitive processing of imagined touch and experienced touch share a common neural substrate (e.g. activate the same population of neurons in similar ways). In this variant, the participant was presented with either objective touch stimuli or instructed to imagine the sensation of being touched. NS was instructed to keep her eyes closed throughout. Objective stimuli were cued to the experimenter with written words that appeared on the monitor. Because the participant’s eyes were closed, the participant did not receive any information about the body part that would be stimulated prior to experiencing the touch. The cue was followed by a one second delay and then at the sound of a beep (the “go” signal), rubs at 1Hz were presented with a metallic probe to either the left or right cheeks, shoulders, or hands. During imagined touch trials, an auditory cue was presented to NS instructing her to imagine being touched on her right cheek, shoulder, or hand. The auditory cue consisted of a voice recording of the words “cheek”, “hand”, or “shoulder” with cue duration of approximately 0.5 seconds. After a one second delay, at the sound of the beep, NS imagined touch to the cued body part. We asked the participant to imagine the sensations as alternating 1Hz rubbing motions similar to what she experienced during objective touch trials. A null condition (without objective or imagined touch), not preceded by an auditory cue was used to establish a baseline neural response. Data were recorded on eight separate days. Eight trials of each condition were performed on each testing day. In total, we recorded from 838 sorted units.

### Quantification and statistical analysis

In the analysis of data from the various task paradigms used in this study, we utilized several statistical methods. Some were specific for certain tasks, but others were applicable to multiple sets of data from the different paradigms. For ease of reference, we have described all methods together in this section. Where necessary, we provide specific examples from tasks to help illustrate their use in our analysis.

#### Linear analysis

In order to determine whether a neuron was tuned (i.e., modulated by a specific condition), we fit a linear model to its firing rate during both the stimulus presentation phase and a baseline time window. The neuronal response during the stimulation phase window was summarized as the mean firing rate computed between 0.5 and 2.5 seconds after stimulus presentation onset. The starting time of 0.5 seconds was chosen to minimize the influence of variable experimenter delay in presenting the stimulus. The baseline response was summarized as the mean firing rate during the 1.5 second window before the stimulus presentation cue. Each unit’s firing rate was modeled as a function of condition indicator variables as:

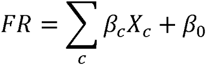

where *FR* is the firing rate, *X*_*c*_ is the vector indicator variable for condition *c*,, *β*_*C*_ is the estimated scalar weighting coefficient for condition *c*, and, *β*_0_ is a constant offset term. In this model, the beta coefficients represent the expected firing rate changes from baseline for each condition. For each condition, the indicator variable is a vector of binary values in which each element is 1 if the corresponding data point at that index is of the same condition type, and 0 if the data point is of a different condition type. All baseline samples were also assigned a 0, effectively pooling together baseline data independent of condition. A unit is considered tuned to a condition if the t-statistic for the beta coefficient associated with the condition is significant (*p*<0.05, false discovery rate (FDR) corrected for multiple comparisons).

#### Discriminability index (DI)

We wished to derive a measure that quantifies how well neural activity can be discriminated from baseline (e.g., pre-stimulus) activity. In other words, we wanted to capture how “well” or how “strongly” a specific sensation (experienced or imagined) is encoded. We developed a cross-validated discriminability index. As with the linear analysis described above, neuronal activity was summarized as the mean firing rate during the stimulation phase window, defined as 0.5 to 2.5 seconds after the onset of stimulus presentation. Baseline phase activity was summarized as the mean firing rate during the 1.5 second window before the stimulus onset presentation cue. The mean activity of all neurons in the population, per condition, was concatenated to form a vector, denoted by *A*. The mean activity of all neurons during the baseline phase was similarly concatenated to form a vector, denoted by *B*. Next, a non-dimensional sensitivity index was computed as:

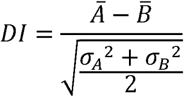

Where *Ā* is the mean of the firing rate vector 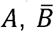 is the mean of the firing rate vector *B,σ*_*A*_ is the standard deviation of the vector *A*, and, *σ*_*B*_ is the standard deviation of the vector.

#### Population Correlation

We used correlation to compare the population neural representations of various tested conditions (stimulus presentations) against each other in a pairwise fashion. Correlation was chosen over alternative distance metrics (such as Mahlanobis or Euclidean distance) because it provides an intuitive metric of similarity that is robust to gross changes in baseline neural activity across the entire neural population. Results were qualitatively similar for alternate distance measures (specifically Mahlanobis distance). In performing correlation analyses, we quantified the neural representations as a vector of firing rates, one vector for each condition with each vector element summarizing the response of an individual unit. As before, neural activity was summarized as the mean firing rate during the stimulation phase window, defined as 0.5 to 2.5 seconds after the onset of stimulus presentation. Firing rate vectors were constructed by averaging the responses across 50-50 splits of trial repetitions. The mean responses across different splits were correlated within and across conditions (e.g. across stimulations of different sensory fields), then the splits were regenerated, and the correlation computed 250 times. The across condition correlations measured similarity between population responses for different sensory fields, answering the question - are the tactile sensations similar or dissimilar from the perspective of the recorded neural population? The within condition correlations assist in our interpretation of the across format correlations by allowing us to quantify the theoretical maxima of the similarity measure (e.g. if the within condition correlation is measured at 0.6, then an across condition of 0.6 suggests identical neural representations.)

To test whether the difference between any pair of conditions was statistically significant, we used a shuffle test applied to the correlations computed over the 250 random splits. To illustrate, in **Figure 5E** we applied this analysis to test whether the correlation between experienced and imagined cheek touch was significantly different from that of experienced cheek touch and imagined shoulder touch. The true difference in the correlations was computed as the difference in the mean correlations between experienced and imagined cheek touches (over the 250 splits) and the mean of the correlations between experienced cheek touch and imagined shoulder touches. We then randomly shuffled the two distributions together (2000 times) and computed the difference in the mean correlations for each shuffle. The distribution of shuffled differences served as the null distribution, against which we compared the true difference to determine significance.

#### Decode analysis (confusion matrix)

Classification was performed using linear discriminant analysis with the following assumptions: one, the prior probability across tested task epochs was uniform; two, the conditional probability distribution of each unit on any epoch was normal; three, only the mean firing rates differ for unit activity during each epoch (covariance of the normal distributions are the same for each); four, firing rates for each input are independent (covariance of the normal distribution is diagonal). The classifier took as input a matrix of average firing rates for all sorted units. The analysis was not limited to significantly modulated units to avoid “peeking” effects. Classification performance is reported as prediction accuracy of a stratified leave-one-out cross-validation analysis. The analysis was performed independently for each recording session and results were then averaged across days.

#### Extent of bilaterality (and generalizability of laterality) in neural representations

The purpose of this analysis was to 1) assess the degree to which tactile information is bilaterally encoded, and to 2) assess the generalizability or similarity in representation for each side to the other (i.e., whether the right and left sides of the body are coded in a similar manner). Only units demonstrating selectivity, that is, differential coding for at least one segment of the body were included in this analysis.

To address the former, for each neuron, we computed the cross-validated coefficient of determination (R^2^_within_) to measure how well a neuron’s firing rate could be explained by the responses to the sensory fields. The R^2^_within_ metric was computed separately for responses to the left (ipsilateral) side and the right (contralateral) side of the body and compared to determine whether the population encoded representations for one body side more robustly than the other.

To address the latter, for each neuron, we computed a regression model using neural data from the ipsilateral side of the body and predicted neural responses for the contralateral side of the body (and vice versa). Predicted responses were compared against true responses to compute a generalization R^2^_across_ metric. This generalization R^2^_across_ was then compared against the cross-validated metric (R^2^_within_) to determine how similar sensory fields were encoded across the left and right sides of the body at the single unit level.

#### Neuronal specificity index

The general formula we used to evaluate the degree of specificity (or specificity index) was:

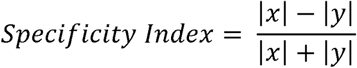

In this formula, x and y are computed on a unit-by-unit basis. The specificity index ranges from −1 to 1, where 1 indicates x ≫ y, and −1 indicates x ≪ y. A value around 0 indicates x ≅ y.

The purpose of computing a specificity index was to quantify the degree to which a neuron was tuned to represent information pertaining to one side of the body over the other. We computed, pairwise, three sets of specificity indices, namely for (1) the cross-validated R^2^_within_ for the right side *x* to the cross-validated R^2^_within_ for the left side y (**Figure 2B**); (2) the cross-validated R^2^_within_ for the left side x to the generalization metric R^2^_across_ when interpreting neural activity from the right side with a left sided prior *y* (**Figure 2D**); and (3) the cross-validated R^2^_within_ for the right side *x* to the generalization metric R^2^_across_ obtained when interpreting neural activity from the left side with a right sided prior y (**Figure 2F**). The *x* and *y* values correspond to those shown in **Figures 2A, 2C, and 2E**, respectively. The specificity index measures a normalized distance from the identity line in each of those figures, with values at −1 corresponding to points above the identity line, and values at +1 corresponding to points below the identity line. A value of zero represents the identity line.

#### Response latency

We quantified the neural response latency to touch stimuli at the level of the neural population. Prior to the analysis, trials were aligned by touch onset as detected by the capacitive touch sensor (ground truth).

Latency was estimated using a sliding window decode analysis: decode performance was computed with k-fold cross-validation of a linear classifier trained over a sliding window through time (linear discriminant analysis with equal diagonal covariance matrices). For visualization purposes, accuracy was computed on data stepped in 50 ms windows and smoothed with a 150 ms full-width at half maximum truncated gaussian smoothing window. For latency calculation, accuracy was computed on data stepped in 2 ms windows and smoothed with a 20 ms full-width at half maximum truncated gaussian smoothing window. For each time window of the latency calculation, significant classification performance was determined when true cross-validated classification was greater than 95% of values of an empirical null distribution of classification accuracies generated by randomly shuffling labels (250 shuffles). Latency is reported as the first window with significant classification. Stimulus conditions were separated by body side (left vs right) and were tested with independent decoders for each body side.

#### Receptive field size

To evaluate the size of touch receptive fields, we characterized the response patterns of individual neurons to tactile stimuli delivered to each of nine points along the subject’s face and neck. Because units mostly demonstrated a single preferred stimulation site, we first identified, for each unit, its preferred point of stimulus delivery as the point associated with the largest firing rate. Next, we examined its response to delivering stimuli to the other points. To estimate the average size of a neuronal receptive field as a function of its preferred point of stimulus delivery, we fit a Gaussian model to the average responses grouped by the preference of the neuron (i.e., data in **Figure 4F**). The Gaussian model had three free parameters, and was defined as:

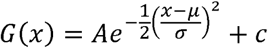

Here, *A* is the amplitude of the Gaussian, is the standard deviation, and is the constant offset term. µ is the mean/center of the Gaussian and was fixed at the preferred point. A separate model (with the appropriate value of µ) was fit to each of the response groups. The full width at half maximum (FWHM) was used to describe the receptive field size.

#### Temporal dynamics of population activity during tactile imagery task: within category

This description relates to **Figure 6B**. We performed a sliding-window classification analysis to quantify the strength and temporal dynamics of population coding in the tactile imagery task. In this task, the participant heard an auditory cue specifying a body part (“cheek”, “hand”, or “shoulder”) that lasted approximately 0.5 seconds, followed by an approximately two second delay, and finally a beep instructing the participant to initiate imagining a touch sensation at the cued body part. This task could engage at least three cognitive processes: 1) semantic processing of the cue; 2) preparation/anticipation for imagery; 3) imagined touch per se. We used a dynamic classification analysis to understand how the neural population evolved through the course of the trial to determine whether the population was best described as mediating a single cognitive processes or multiple cognitive processes. In brief, the analysis consisted of creating a dataset that consisted of the population response measured in small temporal windows throughout the course of the trial. We trained a classifier separately on each temporal window and applied each classifier to both temporal windows. In this way we can measure how information about the cued stimulus evolves in time (e.g. does there exist neural coding during the delay portion of the trial, and, if so, does the neural coding during the delay match neural coding during active imagery). Classification was performed using linear discriminant analysis as described above. We used cross-validation to ensure that training and predicting on the same time window was directly comparable to training on one window and testing on an alternate time window; in other words, we were careful to ensure that accuracy across all comparisons reflects generalization accuracy using the same amount of training and test data. Classifiers were trained and tested on neural responses to the three imagery conditions: cheek, hand, and shoulder. Population response activity for each time window was computed as the average neural response within a 500 ms window, stepped at 100 ms intervals. Window onsets started at −700ms seconds relative to auditory cue onset (cue-delay epoch) with the final window chosen 3.5 seconds after the beep (onset of the imagery epoch). Classification was performed on all sorted units acquired within a single session. Mean and bootstrapped 95% confidence intervals were computed for each time bin from the cross-validated accuracy values computed across sessions.

We believe that this technique, by helping us to understand when information appears and how information compares across task phases, provides a valuable approach to understanding how population activity relates to the underlying cognitive processes. For example, if neural decoding reaches significance only after the go cue, neural activity would be inconsistent with semantic or anticipatory processing. Alternatively, if neural processing begins with the cue, and the same pattern of neural activity is maintained throughout the trial, with no changes during the active imagery phase, then the data would be inconsistent with processing imagined touch per se.

The classification analysis described above was used to measure general similarity of the population response to the tested conditions across time. However, to explicitly test whether population activity was changing, we used Mahalanobis distance as our measure. This is necessary as classification involves a discretization step that makes the technique relatively insensitive to changes in neural population activity that do not cross decision thresholds. Mahalanobis distance, being a proper distance measure, is a more sensitive measure of change. To illustrate, imagine that a classifier is trained on time point A and tested on time point B. At time point A, the means of the two classes are 0 and 1 respectively and at time point 2 the means are 0 and 4 respectively. All classes are assumed to have equal but negligible variance (e.g. 0.01) in this example. When trained on time point A, the classifier finds a decision boundary at 0.5. with 100% classification accuracy. When tested on time point B, with the same 0.5 decision boundary, the classifier again is 100%. Naively, this could be interpreted as signifying that no change in the underlying data has occurred, even though the mean of the second distribution has shifted.

Separation in neural activity between the cue-delay epoch and the imagery epoch was quantified using a cross-validated Mahalanobis distance computed between the observed neural activity at a time point and a reference (baseline) defined as the neural activity immediately following the presentation of the auditory cue, from .25 to .75 seconds. Distances were measured separately for each of the three conditions and then averaged. The mean and standard error on the mean (SEM) were computed across sessions for the cross-validated distance measures and plotted in **Figure 6C**. Activity during the cue-delay epoch and the go epoch were compared using a rank-sum test of the averaged activity during the phase averaged responses across sessions.

#### Temporal dynamics of single unit activity during tactile imagery task: within category

We wished to understand the behavior of single neurons that led to the temporal dynamics of the population. The temporal dynamics of single unit activity during the imagery task (for the imagined touch conditions only) were quantified using both a cluster analysis (**Figure 7A**), and a principle component analysis (PCA, **Figure 7B**). For both, a sliding-window classification analysis was first performed on each sorted unit from all testing days in the same manner as described above for the population activity, with the exception that classifier took as input a vector of the firing rates for a single unit as opposed to a matrix of the firing rates for all units recorded in a single session. This allowed a quantitative description of the temporal dynamics for each sorted unit. We next restricted neurons to those whose 90^th^ percentile accuracy was at least 50%. This was to ensure only neurons with some degree of significant selectivity were used for the cluster analysis. Next, a cluster analysis was performed on these matrices using K-means clustering and the cosine distance metric (chosen to provide a measure of temporal similarity in neural activity profiles, robust to the decode accuracy itself.) We tested cluster sizes from 2 to 5 clusters and used Bayesian information criteria (BIC) to identify the most parsimonious number of clusters for the observed data. In the second analysis, a principal component analysis was applied to the dynamic classification matrices with individual neurons counting as the independent observations. PCA has become a standard method for describing the behavior of neural populations. Typically, PCA is applied to firing rate measurements of neurons. However, in our case, we were less interested in capturing the main modes of variability with respect to individual conditions, but instead wanted to capture the main modes of variability with respect to the temporal dynamics of information encoding.

#### Temporal dynamics of population activity during tactile imagery task: across category

This description pertains to **Figure 8A**. Time-resolved classification analysis was performed using linear discriminate analysis with assumptions and cross-validation procedures as described for within category decoding above. For this analysis, both the experienced touch condition and the imagined touch condition within the tactile imagery task were used. For the experienced touch category, only stimuli to the right cheek, shoulder and hand were used in the analysis; neural activity from the left was not used, to try and match the conditions for the imagined touch conditions in which only right cheek, shoulder, and hand were tested. Classifiers were trained within category and applied to either itself or the other category during each fold. Predictions across folds of the cross-validation procedure were used to compute decode accuracy. This enables us to understand how well the neural representation of the two categories generalize to each other, as well as how well neural representations generalize from one epoch (cue-delay) to another (stimulus: imagined or experienced touch).

## Acknowledgments

The authors would like to thank subject N.S. for participating in the studies, Viktor Shcherbatyuk for technical assistance, and Kelsie Pejsa for administrative and regulatory assistance.

## REFERENCES

1. Buneo CA, Andersen RA. The posterior parietal cortex: sensorimotor interface for the planning and online control of visually guided movements. Neuropsychologia. 2006;44(13):2594–606.

2. Whitlock JR. Posterior parietal cortex. Curr Biol. 2017;27(14):R691–R5.

3. Aflalo T, Kellis S, Klaes C, Lee B, Shi Y, Pejsa K, et al. Decoding motor imagery from the posterior parietal cortex of a tetraplegic human. Science. 2015;348(6237):906–10.

4. Rutishauser U, Aflalo T, Rosario ER, Pouratian N, Andersen RA. Single-Neuron Representation of Memory Strength and Recognition Confidence in Left Human Posterior Parietal Cortex. Neuron. 2018;97(1):209–20 e3.

5. Zhang CY, Aflalo T, Revechkis B, Rosario E, Ouellette D, Pouratian N, et al. Preservation of Partially Mixed Selectivity in Human Posterior Parietal Cortex across Changes in Task Context. eNeuro. 2020;7(2).

6. Zhang CY, Aflalo T, Revechkis B, Rosario ER, Ouellette D, Pouratian N, et al. Partially Mixed Selectivity in Human Posterior Parietal Association Cortex. Neuron. 2017;95(3):697–708 e4.

7. Aflalo T, Zhang, C. Y., Rosario, E., Pouratian, N., Orban, G. A., Andersen, R. A. A shared neural substrate for action verbs and observed actions in human posterior parietal cortex. BioRxiv. 2020.

8. Avillac M, Ben Hamed S, Duhamel JR. Multisensory integration in the ventral intraparietal area of the macaque monkey. J Neurosci. 2007;27(8):1922–32.

9. Graziano MS. Where is my arm? The relative role of vision and proprioception in the neuronal representation of limb position. Proc Natl Acad Sci U S A. 1999;96(18):10418–21.

10. Graziano MS. A system of multimodal areas in the primate brain. Neuron. 2001;29(1):4–6.

11. Graziano MS, Cooke DF, Taylor CS. Coding the location of the arm by sight. Science. 2000;290(5497):1782–6.

12. Graziano MS, Gross CG. A bimodal map of space: somatosensory receptive fields in the macaque putamen with corresponding visual receptive fields. Exp Brain Res. 1993;97(1):96–109.

13. Holmes NP, Spence C. The body schema and the multisensory representation(s) of peripersonal space. Cogn Process. 2004;5(2):94–105.

14. Hwang EJ, Hauschild M, Wilke M, Andersen RA. Spatial and temporal eye-hand coordination relies on the parietal reach region. J Neurosci. 2014;34(38):12884–92.

15. Seelke AM, Padberg JJ, Disbrow E, Purnell SM, Recanzone G, Krubitzer L. Topographic Maps within Brodmann’s Area 5 of macaque monkeys. Cereb Cortex. 2012;22(8):1834–50.

16. Sereno MI, Huang RS. Multisensory maps in parietal cortex. Curr Opin Neurobiol. 2014;24(1):39–46.

17. Binkofski F, Dohle C, Posse S, Stephan KM, Hefter H, Seitz RJ, et al. Human anterior intraparietal area subserves prehension: a combined lesion and functional MRI activation study. Neurology. 1998;50(5):1253–9.

18. Brang D, Taich ZJ, Hillyard SA, Grabowecky M, Ramachandran VS. Parietal connectivity mediates multisensory facilitation. Neuroimage. 2013;78:396–401.

19. Pasalar S, Ro T, Beauchamp MS. TMS of posterior parietal cortex disrupts visual tactile multisensory integration. Eur J Neurosci. 2010;31(10):1783–90.

20. Chan AW, Baker CI. Seeing is not feeling: posterior parietal but not somatosensory cortex engagement during touch observation. J Neurosci. 2015;35(4):1468–80.

21. Keysers C, Wicker B, Gazzola V, Anton JL, Fogassi L, Gallese V. A touching sight: SII/PV activation during the observation and experience of touch. Neuron. 2004;42(2):335–46.

22. Blakemore SJ, Bristow D, Bird G, Frith C, Ward J. Somatosensory activations during the observation of touch and a case of vision-touch synaesthesia. Brain. 2005;128(Pt 7):1571–83.

23. Lucas MV, Anderson LC, Bolling DZ, Pelphrey KA, Kaiser MD. Dissociating the Neural Correlates of Experiencing and Imagining Affective Touch. Cereb Cortex. 2015;25(9):2623–30.

24. Sun HC, Welchman AE, Chang DHF, Di Luca M. Look but don’t touch: Visual cues to surface structure drive somatosensory cortex. Neuroimage. 2016;128:353–61.

25. Keysers C, Gazzola V. Expanding the mirror: vicarious activity for actions, emotions, and sensations. Curr Opin Neurobiol. 2009;19(6):666–71.

26. Sakellaridi S, Christopoulos VN, Aflalo T, Pejsa KW, Rosario ER, Ouellette D, et al. Intrinsic Variable Learning for Brain-Machine Interface Control by Human Anterior Intraparietal Cortex. Neuron. 2019;102(3):694–705 e3.

27. Rosenbaum DA. The movement precuing technique: assumptions, applications and extensions. In: Magill RA, editor. Memory and Control of Action. Amsterdam 1983. p. 231–74.

28. Lecas JC, Requin J, Anger C, Vitton N. Changes in neuronal activity of the monkey precentral cortex during preparation for movement. J Neurophysiol. 1986;56(6):1680–702.

29. Ames KC, Ryu SI, Shenoy KV. Simultaneous motor preparation and execution in a last-moment reach correction task. Nat Commun. 2019;10(1):2718.

30. Kaas JH. What, if anything, is SI? Organization of first somatosensory area of cortex. Physiol Rev. 1983;63(1):206–31.

31. Kaas JH, Nelson RJ, Sur M, Lin CS, Merzenich MM. Multiple representations of the body within the primary somatosensory cortex of primates. Science. 1979;204(4392):521–3.

32. Sur M, Merzenich MM, Kaas JH. Magnification, receptive-field area, and “hypercolumn” size in areas 3b and 1 of somatosensory cortex in owl monkeys. J Neurophysiol. 1980;44(2):295–311.

33. Ferezou I, Haiss F, Gentet LJ, Aronoff R, Weber B, Petersen CC. Spatiotemporal dynamics of cortical sensorimotor integration in behaving mice. Neuron. 2007;56(5):907–23.

34. Geyer S, Schormann T, Mohlberg H, Zilles K. Areas 3a, 3b, and 1 of human primary somatosensory cortex. Part 2. Spatial normalization to standard anatomical space. Neuroimage. 2000;11(6 Pt 1):684–96.

35. Iwamura Y. Hierarchical somatosensory processing. Curr Opin Neurobiol. 1998;8(4):522–8.

36. Jiang W, Tremblay F, Chapman CE. Neuronal encoding of texture changes in the primary and the secondary somatosensory cortical areas of monkeys during passive texture discrimination. J Neurophysiol. 1997;77(3):1656–62.

37. Ruben J, Schwiemann J, Deuchert M, Meyer R, Krause T, Curio G, et al. Somatotopic organization of human secondary somatosensory cortex. Cereb Cortex. 2001;11(5):463–73.

38. Schnitzler A, Salmelin R, Salenius S, Jousmaki V, Hari R. Tactile information from the human hand reaches the ipsilateral primary somatosensory cortex. Neurosci Lett. 1995;200(1):25–8.

39. Tame L, Braun C, Holmes NP, Farne A, Pavani F. Bilateral representations of touch in the primary somatosensory cortex. Cogn Neuropsychol. 2016;33(1-2):48–66.

40. Tame L, Pavani F, Papadelis C, Farne A, Braun C. Early integration of bilateral touch in the primary somatosensory cortex. Hum Brain Mapp. 2015;36(4):1506–23.

41. Zhu Z, Disbrow EA, Zumer JM, McGonigle DJ, Nagarajan SS. Spatiotemporal integration of tactile information in human somatosensory cortex. BMC Neurosci. 2007;8:21.

42. Burton H, Sinclair RJ. Second somatosensory cortical area in macaque monkeys. I. Neuronal responses to controlled, punctate indentations of glabrous skin on the hand. Brain Res. 1990;520(1-2):262–71.

43. de Lafuente V, Romo R. Neural correlate of subjective sensory experience gradually builds up across cortical areas. Proc Natl Acad Sci U S A. 2006;103(39):14266–71.

44. Dykes RW. Parallel processing of somatosensory information: a theory. Brain Res. 1983;287(1):47–115.

45. Garraghty PE, Florence SL, Kaas JH. Ablations of areas 3a and 3b of monkey somatosensory cortex abolish cutaneous responsivity in area 1. Brain Res. 1990;528(1):165–9.

46. Pei YC, Denchev PV, Hsiao SS, Craig JC, Bensmaia SJ. Convergence of submodality-specific input onto neurons in primary somatosensory cortex. J Neurophysiol. 2009;102(3):1843–53.

47. Pons TP, Garraghty PE, Friedman DP, Mishkin M. Physiological evidence for serial processing in somatosensory cortex. Science. 1987;237(4813):417–20.

48. Saal HP, Harvey MA, Bensmaia SJ. Rate and timing of cortical responses driven by separate sensory channels. Elife. 2015;4:e10450.

49. Sakata H, Takaoka Y, Kawarasaki A, Shibutani H. Somatosensory properties of neurons in the superior parietal cortex (area 5) of the rhesus monkey. Brain Res. 1973;64:85–102.

50. Soso MJ, Fetz EE. Responses of identified cells in postcentral cortex of awake monkeys during comparable active and passive joint movements. J Neurophysiol. 1980;43(4):1090–110.

51. Iwamura Y. Bilateral receptive field neurons and callosal connections in the somatosensory cortex. Philos Trans R Soc Lond B Biol Sci. 2000;355(1394):267–73.

52. Iwamura Y, Iriki A, Tanaka M. Bilateral hand representation in the postcentral somatosensory cortex. Nature. 1994;369(6481):554–6.

53. Iwamura Y, Tanaka M, Iriki A, Taoka M, Toda T. Processing of tactile and kinesthetic signals from bilateral sides of the body in the postcentral gyrus of awake monkeys. Behav Brain Res. 2002;135(1-2):185–90.

54. Taoka M, Toda T, Iriki A, Tanaka M, Iwamura Y. Bilateral receptive field neurons in the hindlimb region of the postcentral somatosensory cortex in awake macaque monkeys. Exp Brain Res. 2000;134(2):139–46.

55. Chand P, Jain N. Intracortical and Thalamocortical Connections of the Hand and Face Representations in Somatosensory Area 3b of Macaque Monkeys and Effects of Chronic Spinal Cord Injuries. J Neurosci. 2015;35(39):13475–86.

56. Chang SW, Dickinson AR, Snyder LH. Limb-specific representation for reaching in the posterior parietal cortex. J Neurosci. 2008;28(24):6128–40.

57. Taoka M, Toda T, Iwamura Y. Representation of the midline trunk, bilateral arms, and shoulders in the monkey postcentral somatosensory cortex. Exp Brain Res. 1998;123(3):315–22.

58. Beurze SM, de Lange FP, Toni I, Medendorp WP. Integration of target and effector information in the human brain during reach planning. J Neurophysiol. 2007;97(1):188–99.

59. Friedman DP, Murray EA. Thalamic connectivity of the second somatosensory area and neighboring somatosensory fields of the lateral sulcus of the macaque. J Comp Neurol. 1986;252(3):348–73.

60. Berger CC, Ehrsson HH. The fusion of mental imagery and sensation in the temporal association cortex. J Neurosci. 2014;34(41):13684–92.

61. Wise NJ, Frangos E, Komisaruk BR. Activation of sensory cortex by imagined genital stimulation: an fMRI analysis. Socioaffect Neurosci Psychol. 2016;6:31481.

62. Gentile G, Petkova VI, Ehrsson HH. Integration of visual and tactile signals from the hand in the human brain: an FMRI study. J Neurophysiol. 2011;105(2):910–22.

63. Keysers C, Gazzola V. Dissociating the ability and propensity for empathy. Trends Cogn Sci. 2014;18(4):163–6.

64. Klaes C, Kellis S, Aflalo T, Lee B, Pejsa K, Shanfield K, et al. Hand Shape Representations in the Human Posterior Parietal Cortex. J Neurosci. 2015;35(46):15466–76.

65. Yang Y, Dickey MW, Fiez J, Murphy B, Mitchell T, Collinger J, et al. Sensorimotor experience and verb-category mapping in human sensory, motor and parietal neurons. Cortex. 2017;92:304–19.

66. Fusi S, Miller EK, Rigotti M. Why neurons mix: high dimensionality for higher cognition. Curr Opin Neurobiol. 2016;37:66–74.

67. Binder JR, Desai RH. The neurobiology of semantic memory. Trends Cogn Sci. 2011;15(11):527–36.

68. Meyer K, Damasio A. Convergence and divergence in a neural architecture for recognition and memory. Trends Neurosci. 2009;32(7):376–82.

69. Lambon Ralph MA, Jefferies E, Patterson K, Rogers TT. The neural and computational bases of semantic cognition. Nature Reviews Neuroscience. 2017;18(1):42–55.

70. Patterson K, Nestor PJ, Rogers TT. Where do you know what you know? The representation of semantic knowledge in the human brain. Nat Rev Neurosci. 2007;8(12):976–87.

71. Miyashita Y. Perirhinal circuits for memory processing. Nat Rev Neurosci. 2019;20(10):577–92.

72. Makin TR, Bensmaia SJ. Stability of Sensory Topographies in Adult Cortex. Trends Cogn Sci. 2017;21(3):195–204.

73. Armenta Salas M, Bashford L, Kellis S, Jafari M, Jo H, Kramer D, et al. Proprioceptive and cutaneous sensations in humans elicited by intracortical microstimulation. Elife. 2018;7.

74. Flesher SN, Collinger JL, Foldes ST, Weiss JM, Downey JE, Tyler-Kabara EC, et al. Intracortical microstimulation of human somatosensory cortex. Sci Transl Med. 2016;8(361):361ra141.

75. Tandon S, Kambi N, Lazar L, Mohammed H, Jain N. Large-scale expansion of the face representation in somatosensory areas of the lateral sulcus after spinal cord injuries in monkeys. J Neurosci. 2009;29(38):12009–19.

76. Mohammed H, Hollis ER, 2nd. Cortical Reorganization of Sensorimotor Systems and the Role of Intracortical Circuits After Spinal Cord Injury. Neurotherapeutics. 2018;15(3):588–603.

77. Brainard DH. The Psychophysics Toolbox. Spatial Vision. 1997;10(4):433–6.

